# Cell wall melanin impedes growth of the *Cryptococcus neoformans* polysaccharide capsule by sequestering calcium

**DOI:** 10.1101/2024.06.20.599928

**Authors:** Rosanna P. Baker, Amy Z. Liu, Arturo Casadevall

## Abstract

*Cryptococcus neoformans* has emerged as a frontrunner among deadly fungal pathogens and is particularly life-threatening for many HIV-infected individuals with compromised immunity. Multiple virulence factors contribute to the growth and survival of *C. neoformans* within the human host, the two most prominent of which are the polysaccharide capsule and melanin. As both of these features are associated with the cell wall, we were interested to explore possible cooperative or competitive interactions between these two virulence factors. Whereas capsule thickness had no effect on the rate at which cells became melanized, build-up of the melanin pigment layer resulted in a concomitant loss of polysaccharide material, leaving melanized cells with significantly thinner capsules than their non-melanized counterparts. When melanin was provided exogenously to cells in a transwell culture system we observed a similar inhibition of capsule growth and maintenance. Our results show that melanin sequesters calcium thereby limiting its availability to form divalent bridges between polysaccharide subunits required for outer capsule assembly. The decreased ability of melanized cells to incorporate exported polysaccharide into the growing capsule correlated with the amount of shed polysaccharide, which could have profound negative impacts on the host immune response.

**Significance Statement:** *Cryptococcus neoformans* is an opportunistic fungal pathogen that presents a significant health risk for immunocompromised individuals. We report an interaction between the two major cryptococcal virulence factors, the polysaccharide capsule and melanin. Melanin impacted the growth and maintenance of the polysaccharide capsule, resulting in loss of capsular material during melanization. Our results suggest that melanin can act as a sink for calcium, thereby limiting its availability to form ionic bridges between polysaccharide chains on the growing surface of the outer capsule. As polysaccharide is continuously exported to support capsule growth, failure of melanized cells to incorporate this material results in a higher concentration of shed polysaccharide in the extracellular milieu, which is expected to interfere with host immunity.

## Introduction

In 2022, *Cryptococcus neoformans* was declared as a ‘critical’ human pathogenic fungus by the World Health Organization because of its ability to cause life-threatening disease (1). Cryptococcosis is more likely to occur in individuals with impaired immunity, such as those with advanced HIV infection, and has a high morbidity and mortality despite therapy with antifungal drugs (2). Cryptococcal disease is notorious for central nervous system involvement where it often presents as meningoencephalitis. Therapy requires long courses of antifungal drug treatment and even among those who respond to therapy recurrences are common (3). There are currently no vaccines available to prevent cryptococcosis in those at high risk of disease.

*C. neoformans* is a species complex (4) that encompasses several species with human pathogenic potential sharing a suite of virulence factors that includes a polysaccharide capsule (5), a laccase that catalyzes the synthesis of melanin (6), urease (7) and several other enzymes (8, 9). Despite genetic differences these species cause similar clinical disease, possibly because they share the same virulence factors and elicit similar inflammatory responses. Each of the cryptococcal virulence factors contributes to the capacity of the fungus to resist clearance by the immune response. The capsule is antiphagocytic, melanin protects against oxidative fluxes and urease has pleiotropic effects on the immune response and survival in macrophages. Cryptococcal virulence factors, believed to have been selected for environmental survival as mechanisms of defense against ameboid predators, also confer survival advantages when fungal cells are attacked by immune cells (10, 11).

Among these virulence factors the capsule and melanin make the greatest contribution to the cryptococcal virulence phenotype (12, 13). Although each of these virulence factors has been extensively studied for their singular contribution to virulence, relatively little work has been done to understand how they interact to promote fungal survival in the environment and the host. Recently, we described that melanization and urease activity were reciprocally regulated through a mechanism whereby increased pH resulting from the conversion of urea into ammonia promoted melanization, while melanization in turn reduced the release of urease containing vesicles (14).

In this study we followed the incidental observation made while studying the contribution of capsule and melanin to buoyant density that melanized cells had smaller capsules than non-melanized cells (15) to understand the relationship between these virulence factors. As the greatest contributors to cryptococcal virulence, the existence of a regulatory relationship between capsule and melanin, if any, could have important implications for understanding cryptococcal virulence because they share similarities and differences in the properties they confer on cryptococci. The capsule is antiphagocytic (16) but also functions to absorb oxygen- and nitrogen-derived radicals produced during the oxidative burst (17). Capsular polysaccharide is also released into infected tissues where it can mediate a variety of deleterious effects on the immune response (18). Melanin confers structural strength to the cell wall (19), while also protecting cryptococcal cells against microbicidal peptides (20) and serves as a sink for absorbing oxidative radicals (21). Exploration of the correlation between melanization and thinner capsules revealed that the effect is mediated by the Ca^2+^ binding properties of melanin that prevent calcium-dependent capsule growth resulting in increased release of polysaccharide, an effect that could have profound consequences in pathogenesis as the released polysaccharide interferes with cellular and immune processes.

## Results

### Melanization is associated with smaller capsules

Wild-type *C. neoformans* strain, KN99*α*, was grown in minimal media in the absence or presence of dopamine for 2 d and the cells were then suspended in India ink to visualize the polysaccharide capsule (Fig. 1A). Cells that had become pigmented by synthesizing melanin from dopamine had an average capsule thickness of 1.5 µm, which was significantly smaller than the average capsule thickness of 4.5 µm measured for non-melanized cells (Fig. 1B, left panel). Melanized cells were also larger, with an average cell radius of 2.6 µm, compared to an average cell radius of 2.3 µm for non-melanized cells (Figure 1B, center panel). Comparison of the capsule/body ratio revealed an average ratio of 0.6 for melanized cells, which was significantly lower than the average ratio of 1 for non-melanized cells (Fig. 1B, right panel). These results are consistent with a prior report of larger cell size and smaller capsules for melanized cells compared to non-melanized cells of the H99 *C. neoformans* strain (15).

**Fig. 1.**
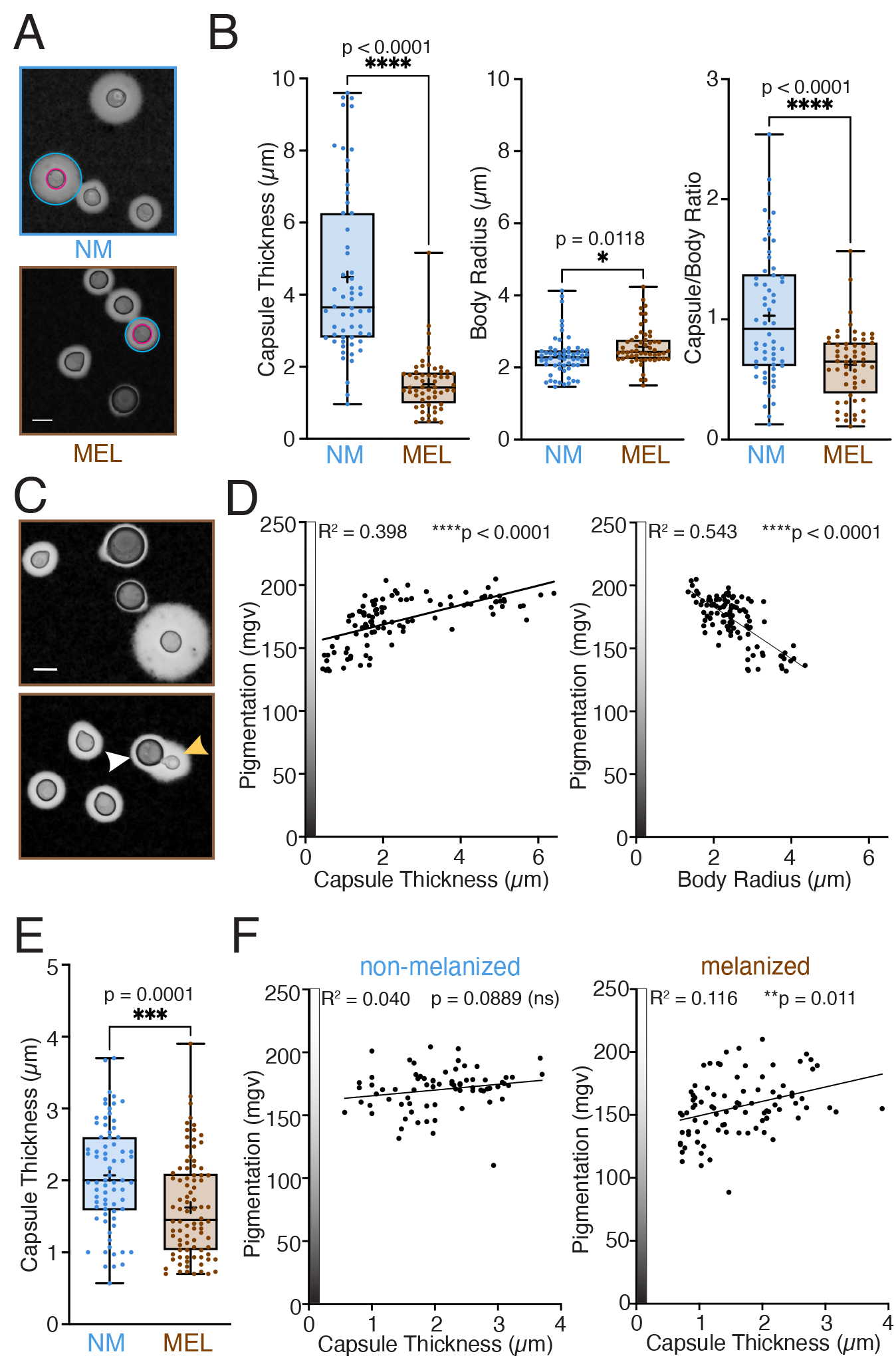
Melanized cells have smaller capsules than non-melanized cells *in vitro* and *in vivo*. (A) India ink microscopy images of non-melanized (NM) and melanized (MEL) cells grown for 2 d in minimal media (MM) in the absence or presence of 1 mM dopamine, respectively. For one cell in each image, the cell body is outlined in magenta and the capsule is outlined in cyan. (B) Compared to NM cells, MEL cells have thinner capsules (left panel), are larger in size (center panel), and have smaller capsule/body ratios (right panel). (C) India ink images of cells grown for 24 h in dopamine-supplemented MM reveals a mixture of heavily pigmented cells with small capsules and pale cells with large capsules (upper panel). The lower panel shows a heavily pigmented mother cell with a thin capsule (white arrow) and pale daughter cell with a thicker capsule (yellow arrow). (D) Increased pigmentation correlates linearly with decreased capsule thickness (left panel) and increased cell size (right panel). (E) Comparison of capsule thickness of cells recovered from mouse lung tissue 24 h after infection with macrophages harboring either NM or MEL cells. (F) Increased pigmentation correlates linearly with decreased capsule size for the melanized (right panel) but not for the non-melanized cells (left panel). Data were analyzed using an unpaired parametric t-test (B and E) or Pearson correlation with two-tailed P value (D and F). Scale bar is 5 µm.

Melanization is a gradual process whereby pigment granules are deposited in concentric layers on the cryptococcal cell wall (22). Microscopic examination of cells growing in dopamine- supplemented media in the first day of culture, when cells are still actively dividing, revealed a mixed population of large, heavily pigmented cells with small capsules and small, pale cells with large capsules (Fig. 2C, upper panel). For melanized cells in the process of budding, the mother cell had a much smaller capsule than the newly budded daughter cell (Fig. 2C, lower panel). For a set of approximately 100 cells in this melanizing culture, we performed capsule and cell body measurements and quantified the mean gray value of each cell to assess the degree of pigmentation.

**Fig. 2.**
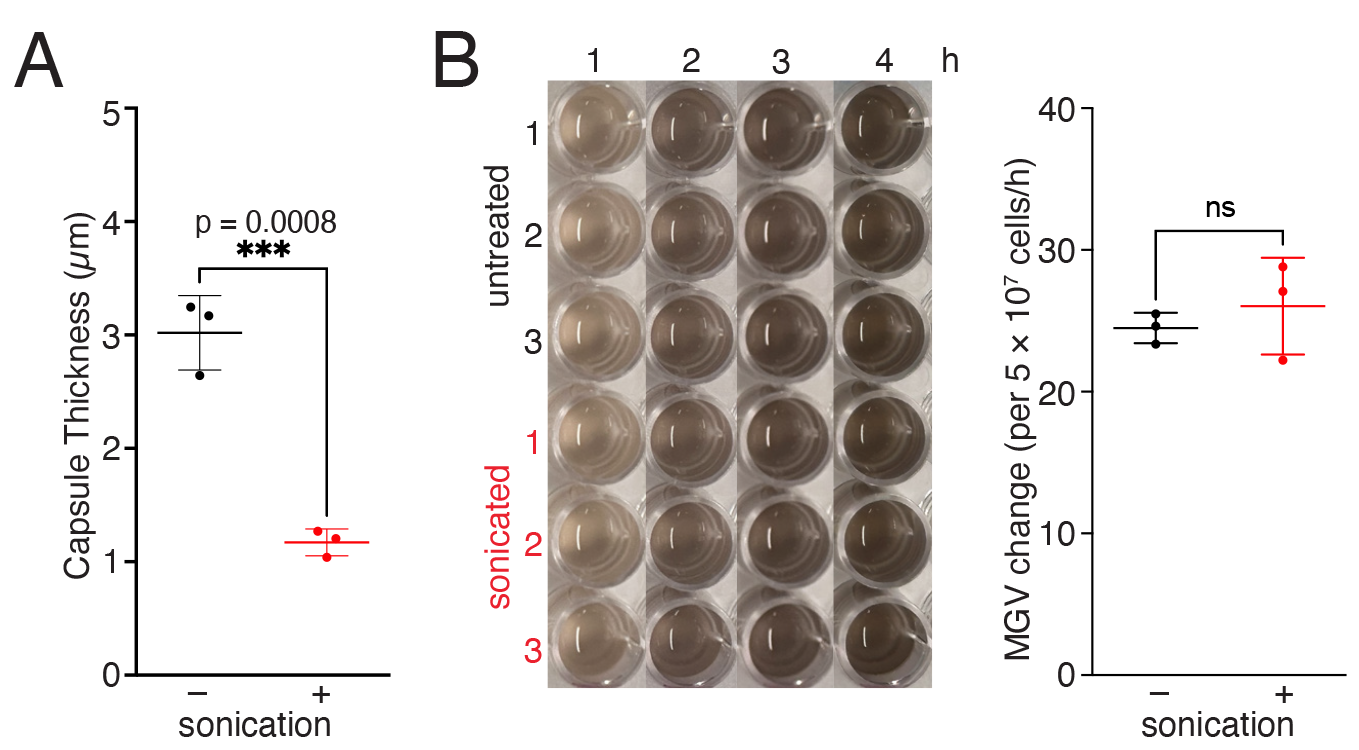
Capsule size does not influence melanin pigment production rate. (A) Capsule thickness measurements for three biological replicate cultures before and after sonication showing a significant loss of capsular material for sonicated cells. (B) Time course of melanin pigment production by untreated and sonicated cells incubated in the presence of 1 mM dopamine (left panel) reveals a comparable rate of change in mean gray value (MGV) for untreated cells with large capsules and sonicated cells with small capsules (right panel). Data were analyzed using an unpaired parametric t-test.

Our analysis revealed a negative correlation between pigmentation and capsule size, with darker cells having smaller capsules than lighter cells (Fig. 1D, left panel). In contrast, cell size correlated positively with pigmentation as darker cells tended to be larger than lighter cells (Fig. 1D, right panel). This observation may reflect cell age since young cells are smaller than older cells and, likewise, would have had less time to build up a layer of melanin on the cell wall.

To investigate whether melanization also impacted capsule size during infection, we prepared India ink suspensions of mouse lung homogenates from mice infected with macrophages harboring either non-melanized or melanized *C. neoformans* cells 24 h post-infection (14). Our analysis revealed significantly smaller capsules for cells recovered from mice infected with melanized compared to non-melanized cells, with average thicknesses of 1.6 and 2.1 µm, respectively (Fig. 1E). Since cell division is likely to have occurred during the 24 h infection window, daughter cells newly derived from melanized mothers are expected to lack pigmentation, as observed *in vitro* (Fig. 1C). We quantified the degree of pigmentation for each cell and plotted it against capsule thickness, which revealed a linear correlation between increased pigmentation and decreased capsule size for cells recovered from mice infected with melanized cells whereas no such correlation was observed with non-melanized cells (Fig. 1F).

### Large capsules do not slow melanin pigment production

Cells with large capsules left untreated or sonicated to significantly reduce capsule size (Fig. 2A) were provided with dopamine and photographed over the course of several hours to monitor melanin pigment production (Fig. 2B, left panel). Quantified rates of mean gray value change over time for three biological replicate untreated and sonicated cultures showed no statistically significant difference (Fig. 2B, right panel). Hence, whereas the presence of cell wall melanin was correlated with decreased capsule size (Fig. 1), we found no evidence for a reciprocal influence of capsule size on melanin deposition.

### Melanization is associated with increased capsular polysaccharide release

Since cells with large capsules readily produced melanin pigment (Fig. 2), we sought to follow the fate of the polysaccharide capsule during the first 24 h of melanization. Within 6 h of growth in the presence of dopamine, average capsule thickness decreased significantly from approximately 4 µm to a steady state of about 2.8 µm (Fig. 3A). The rapid decrease in capsule thickness observed during the first 6h of melanization correlated with a high rate of polysaccharide release into the culture media that subsequently decreased to a significantly lower rate by 24 h (Fig. 3B).

**Fig. 3.**
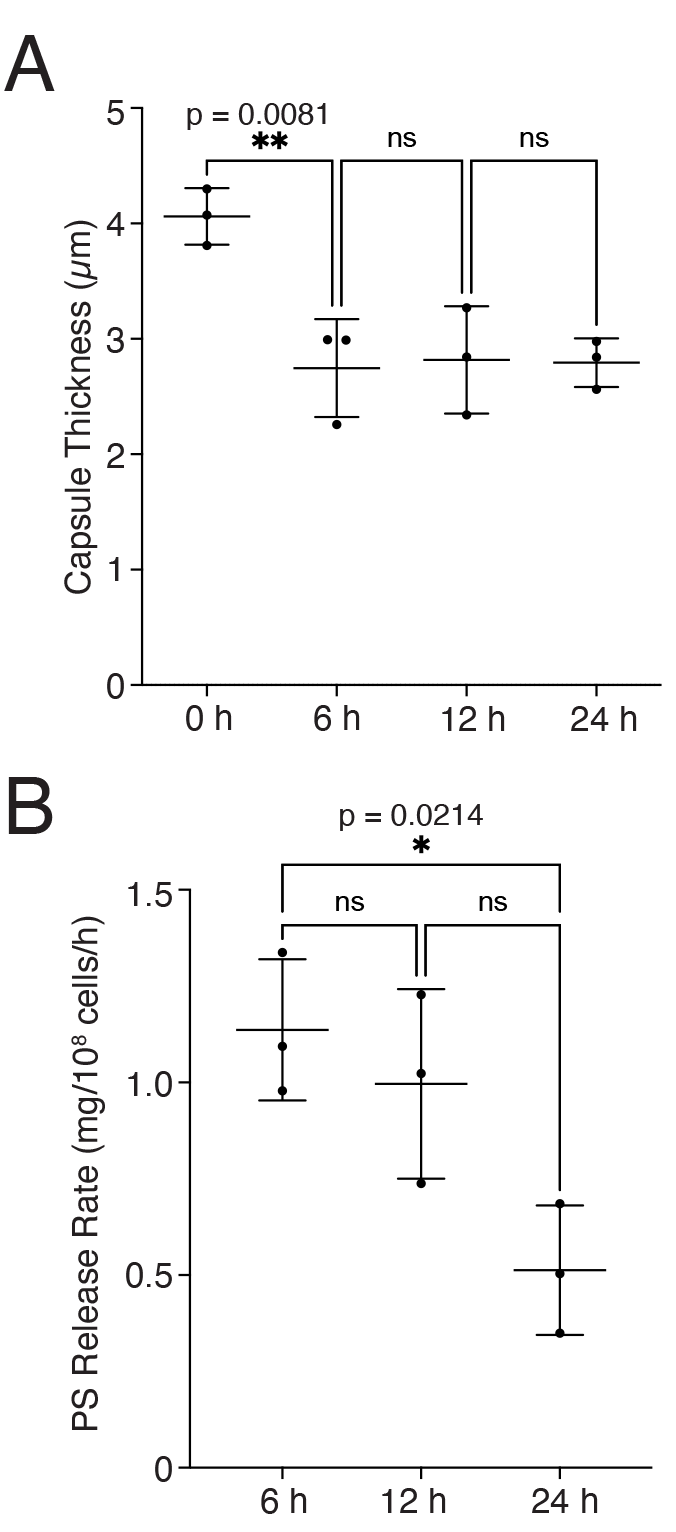
Capsular material is rapidly lost as cells become melanized. (A) Capsule thickness measurements for cells that had been incubated in minimal media for 2 d to induce capsule growth before being sub-cultured in triplicate into media supplemented with dopamine. The average capsule size decreased significantly during the first 6 h of melanization and then remained unchanged for the remaining 18 h. (B) The rate of polysaccharide (PS) release into the culture media measured by GXM ELISA decreased over time during melanization and was significantly lower at 24 h compared to 6 h. Data were analyzed using a one-way analysis of variance (ANOVA) with Tukey’s multiple comparisons test.

### The extent of melanization correlates positively with polysaccharide release

We took advantage of the increased buoyant density of melanized compared to non-melanized cells (15) to enrich for the sub-population of heavily pigmented cells in a melanizing culture using Percoll density gradient centrifugation (Fig. 4A). Following placement of these heavily melanized cells into fresh dopamine-supplemented media, the amount of polysaccharide released into the culture media over time was significantly greater for enriched melanized cultures compared to non-melanized control cultures (Fig. 4B). This observation may explain why cells with a heavy build-up of cell wall melanin manifest smaller average capsule sizes than non-melanized cells (Fig. 1) if the polysaccharide exported extracellularly from melanized cells for capsule construction was lost rather than assimilated into the capsule.

**Fig. 4.**
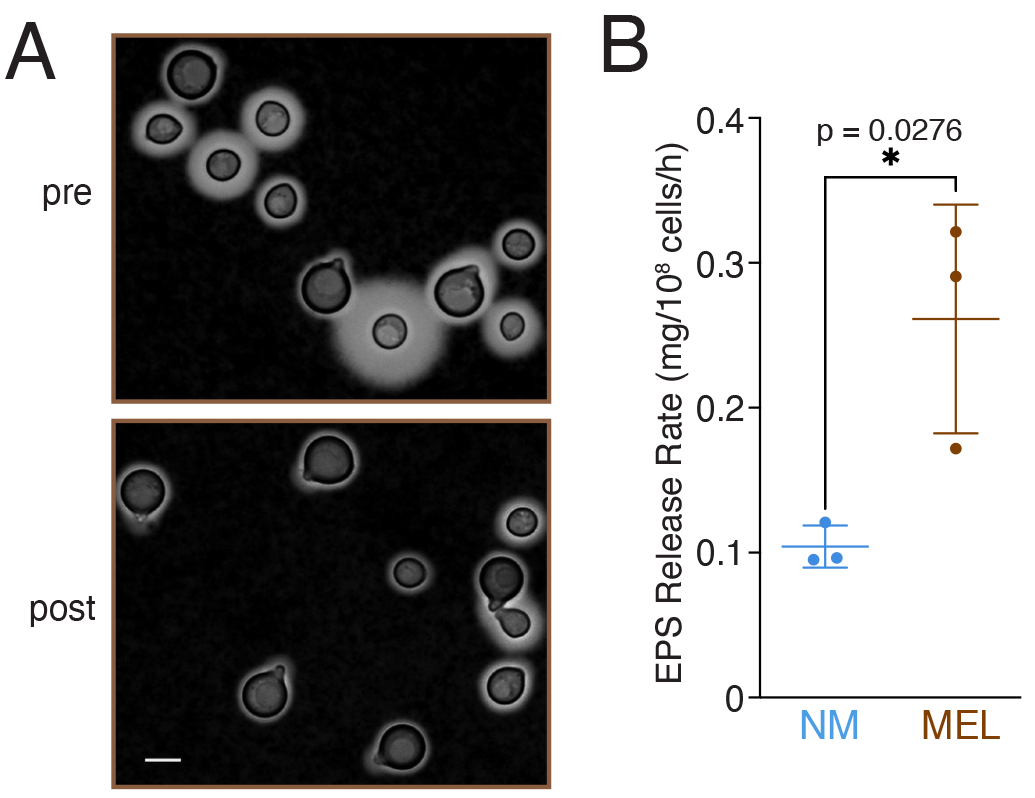
Melanized cells shed more polysaccharide than non-melanized cells. (A) Representative India ink images of cells before (pre) and after (post) enrichment for heavily melanized cells by Percoll density gradient centrifugation. (B) Comparison of exopolysaccharide (EPS) release rate for three biological replicate non-melanized and enriched melanized cultures reveals a significantly higher shedding rate for melanized compared to non-melanized cells. Data were analyzed using an unpaired parametric t-test. Scale bar is 5 µm.

### Capsule removal and regrowth from non-melanized and melanized cells

Polysaccharide was removed from cells in non-melanized and enriched melanized cultures by sonication, as this is the optimal method for removing capsular material without killing cells (23). Capsule thickness for the non-melanized cells decreased substantially from an average of 3 µm to 1 µm while the melanized cells that already had a small average capsule thickness of 1.2 µm decreased further to 0.97 µm (Figure 5A). Cell body and total (combined cell and capsule) radius measurements were used to calculate the capsular volume before and after sonication for non-melanized and melanized cells from three biological replicate cultures. This analysis revealed an approximate 20-fold excess of capsular volume removed from non-melanized compared to melanized cells (Fig. 5B, left panel). By contrast, quantification of the relative yields using polysaccharide ELISA showed only a 2-fold greater mass of polysaccharide removed from non-melanized compared to melanized cells (Fig. 5B, right panel) which implies that melanized capsules are considerably denser.

**Fig. 5.**
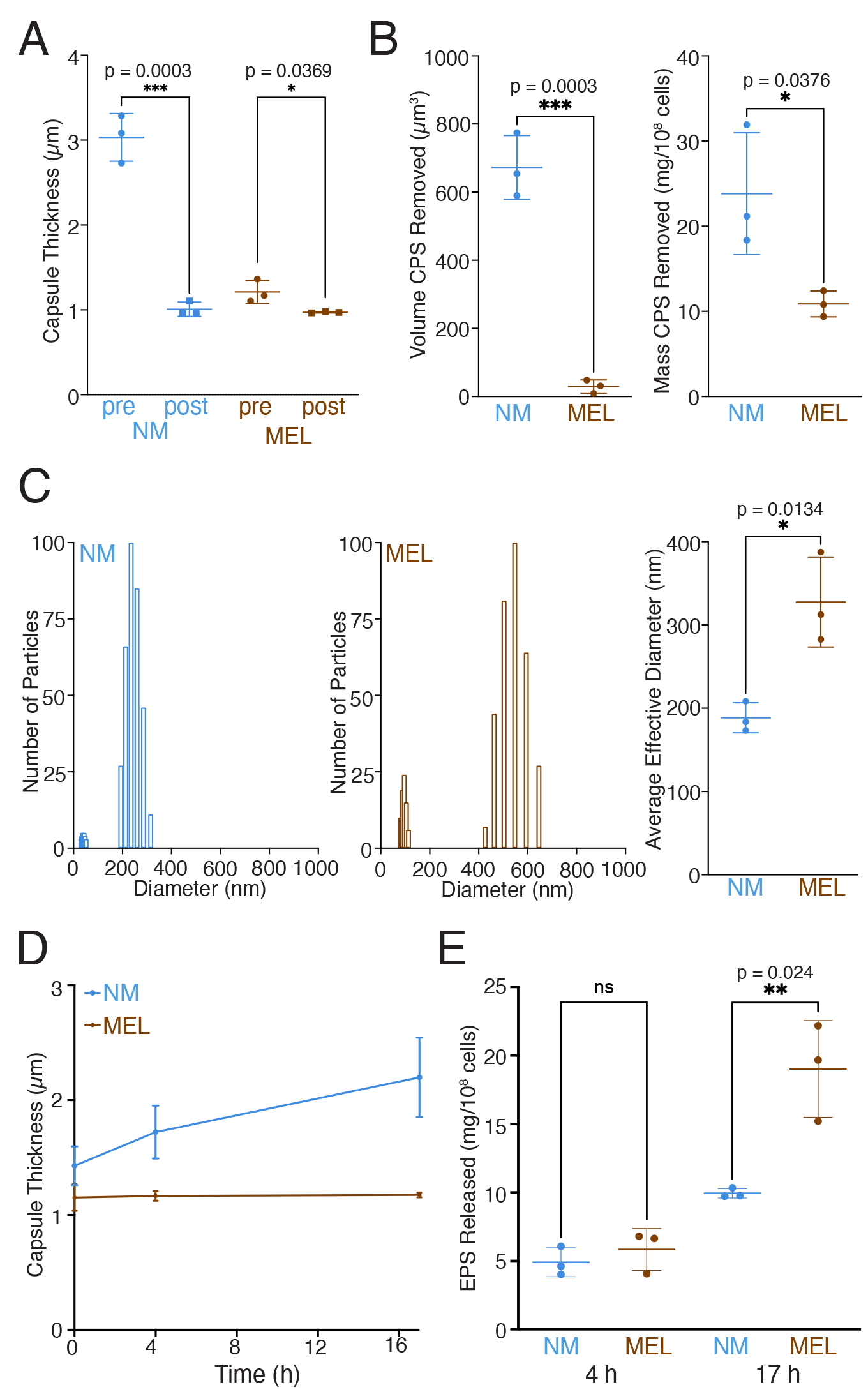
Removal and regrowth of capsules from non-melanized and melanized cells. (A) Capsule thickness measurements for three biological replicate cultures of non-melanized (NM) and enriched melanized (MEL) cells before (pre) and after (post) physical removal of capsular material by sonication. (B) Approximately 20-fold greater volume (left panel) but only 2-fold more mass (right panel) of polysaccharide was removed from NM compared to MEL cells. (C) Dynamic light scattering of polysaccharide collected from NM (left panel) and MEL (center panel) cells reveals a size distribution skewed toward larger particles and an greater average effective diameter (right panel) for MEL compared to NM cells. (D) Time course of capsule regrowth for NM and MEL cells. (E) Significantly more released polysaccharide accumulates in culture media of MEL compared to NM cells during 17 h of capsule regrowth. Statistical significance was determined using an unpaired parametric t-test (A-C) or one-way analysis of variance (ANOVA) with Tukey’s multiple comparisons test (E).

Capsular polysaccharide (CPS) collected from non-melanized and melanized cells was filtered through a 0.8 µm filter to remove any cells that may have been retained in solution after centrifugation and then particle size measurements were conducted using dynamic light scattering. The distribution of particle sizes was shifted toward larger particles for CPS derived from melanized cells, with an average effective diameter of 328 nm, compared to that of 188 nm for non-melanized cells. (Fig. 5C).

Following capsular removal by sonication, non-melanized and melanized cells were sub- cultured into fresh minimal media without and with dopamine, respectively. Capsule thickness measured over the course of 17 h showed significant capsule regrowth only for the non-melanized cells (Fig. 5D). The amount of polysaccharide that had accumulated in culture media was significantly greater for melanized compared to non-melanized cells by 17 h (Fig. 5E). This result suggests that polysaccharide exported extracellularly for capsule assembly more readily accrues onto non-melanized compared to melanized cells to allow capsular regrowth after physical removal.

### Exogenous melanin inhibits capsule growth by depleting calcium

We pondered whether capsule growth was being inhibited by an alteration of cell wall architecture imparted by the melanin pigment layer or by a physical property of the melanin itself. To explore the latter possibility, cells were grown in a 0.4 µm transwell insert with the outer well containing MM or MM supplemented with 5 mg/mL *C. neoformans* melanin ghosts, which are acid resistant shells of cell wall melanin (24), or 3 mg/mL synthetic melanin. Both forms of exogenously added melanin strongly inhibited capsule growth as fewer than 10% of cells grew large capsules compared to approximately 75% for the MM control (Fig. 6A). We extended our analysis to address whether addition of melanin in trans would impact capsule maintenance by culturing cells in MM for 2 d to induce capsule growth prior to sub- culturing into the transwell system. Our results revealed a concentration-dependent effect whereby approximately 75% of cells grown in the presence of 0.5 mg/mL synthetic melanin retained large capsules, which was comparable to the MM control, whereas the percentage of cells with large capsules decreased to approximately 40% and 20% in the presence of 1 mg/mL and 2 mg/mL synthetic melanin, respectively (Fig. 6B).

**Fig. 6.**
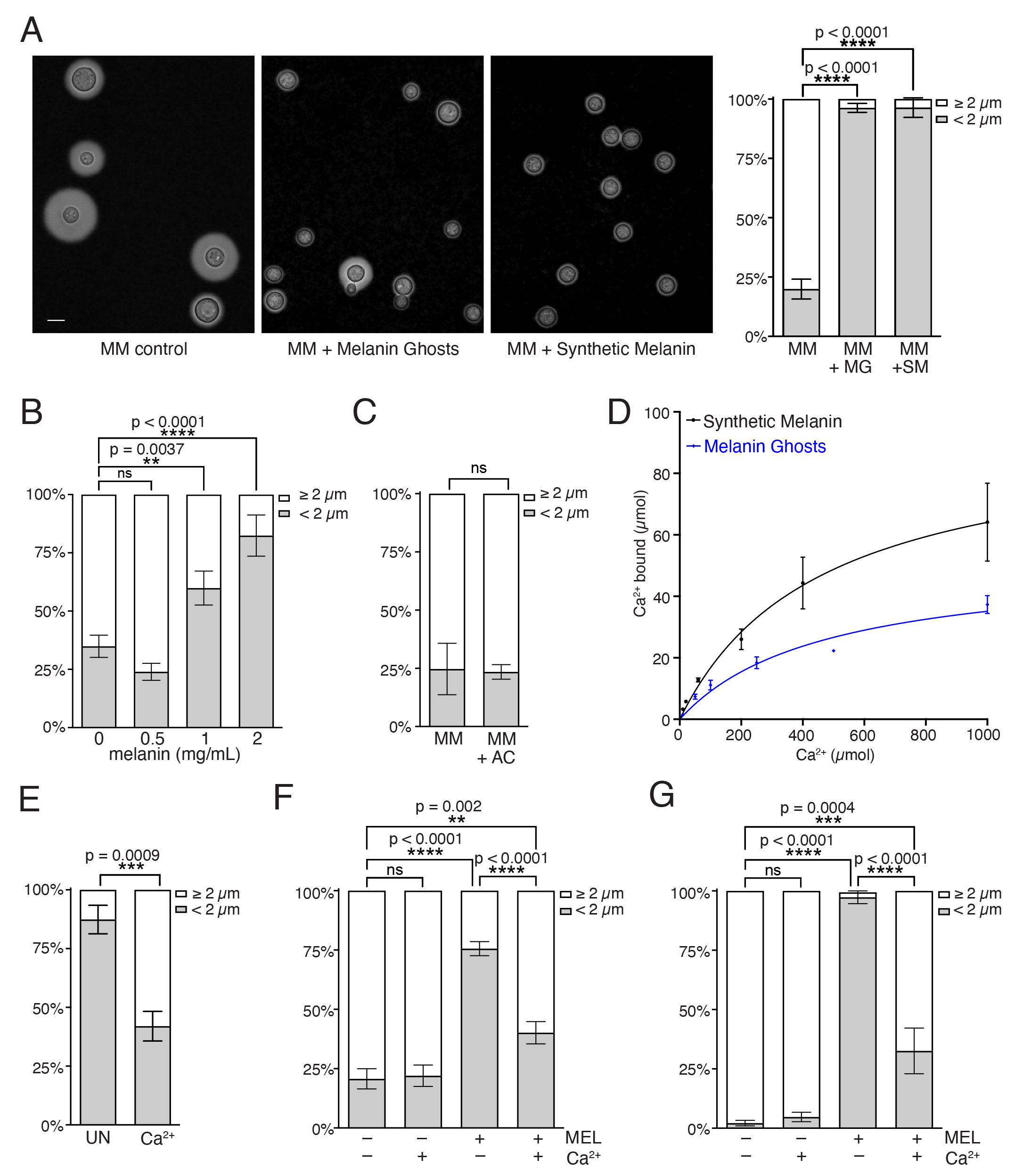
Melanin inhibits capsule growth and maintenance by binding calcium. (A) Representative India ink images (left 3 panels) of cells that had been incubated in transwell culture inserts in wells containing either minimal media only (MM) or MM supplemented with melanin ghosts (MG) or synthetic melanin (SM) and quantified percentages of cells with small (< 2 µm) or large (≥ 2 µm) capsules (right panel). (B) Cells with large capsules from pre-growth in MM were sub-cultured in transwell culture inserts in the presence of the indicated concentration of synthetic melanin and relative percentages of cells with small and large capsules were quantified. (C) Quantified percentages of cells with small (< 2 µm) or large (≥ 2 µm) capsules following growth in a transwell system containing MM or MM supplemented with 5 mg/mL activated charcoal (AC). (D) Saturation Ca^2+^ binding curve plotted as mean and standard deviation for 1 mg synthetic melanin (black) and *C. neoformans* melanin ghosts (blue) (E) Comparison of capsule sizes for cells grown in a transwell system containing MM supplemented with 2 mg/ml synthetic melanin that was untreated (UN) or pre-incubated with 1M calcium chloride (Ca^2+^). (F) Percentages of cells with small and large capsules quantified following growth in a transwell system with MM in the absence (–) or presence (+) of 1 mg/mL synthetic melanin (MEL) or 0.1 mM calcium chloride (Ca^2+^) as indicated. (G) Percentages of cells with small and large capsules quantified following growth in a transwell system with magnesium-free MM in the absence (–) or presence (+) of 1.5 mg/mL synthetic melanin (MEL) or 0.1 mM calcium chloride (Ca^2+^) as indicated. Statistical significance was determined using a one-way analysis of variance (ANOVA) with Tukey’s multiple comparisons test (A, B, F, G) or an unpaired parametric t-test (C, E). Scale bar is 5 µm.

The ability of melanin to inhibit capsule growth and maintenance from a distance suggests that melanin could be binding a soluble factor that is required for the growth of large capsules. We observed no effect on capsule growth when cells were cultured in a transwell containing MM supplemented with 5 mg/mL activated charcoal (Fig. 6C) so it is unlikely that the melanin effect is from the absorption of a small organic molecule. As melanin has a high affinity for a broad range of metal ions (25) and cross-linking of polysaccharide molecules in the *C. neoformans* capsule is coordinated through binding of divalent metal ions (26), sequestration of calcium by melanin provided a plausible explanation for our results. Using a calcium meter, we measured the concentration of calcium remaining in solution after incubation for 1 h with 1 mg of the melanin ghosts or synthetic melanin used in our assays. From these measurements, the concentration of bound calcium was inferred to derive maximal binding capacity values (Bmax) of approximately 52 and 93 µmoles calcium per mg of melanin ghosts and synthetic melanin, respectively (Fig. 6D).

To explore whether calcium binding to melanin was affecting capsule enlargement in the transwell system, we grew cells in the presence of 2 mg/mL synthetic melanin that had been pre-bound to calcium and observed a significantly higher percentage of cells with large capsules in the presence of calcium-saturated compared to untreated melanin (Fig. 6E). We also compared capsule sizes of cells grown in the absence and presence of exogenous melanin in media with or without added calcium. In the absence of melanin, addition of calcium had no effect on capsule size whereas addition of 0.1 mM Ca^2+^ to wells containing 1 mg/mL melanin resulted in a significant increase in the percentage of cells with large capsules, from approximately 24% to 60% (Fig 6F). As GXM binds both Ca^2+^ and Mg^2+^ divalent metal ions (26), we performed a transwell experiment in magnesium-free MM and found that addition of 0.1 mM Ca^2+^ in the absence of Mg^2^ also alleviated the inhibitory effect of melanin on capsule growth (Fig. 6G) In both cases, the inhibitory effect of melanin on capsule growth was partially relieved by added calcium, albeit not fully restored to that observed in the absence of melanin.

## Discussion

The cell wall of *C. neoformans* is the site of assembly for a polysaccharide capsule and melanin pigment layer, both of which make major contributions to survival in the environment and protection from host defense mechanisms during infection. As such, these two virulence factors are key targets in the development of potential prevention and treatment strategies, including polysaccharide-based vaccines (27, 28), melanin inhibitors (29), and antibodies to melanin (30). Having recently discovered an extra level of complexity in the study of *C. neoformans* virulence whereby two individual factors, melanin and urease, interacted to produce unexpected outcomes during infection (14), we sought to explore possible interactions between melanin and the polysaccharide capsule. Whereas no correlation was found between capsule size and the rate of melanization, a negative correlation between melanization and capsule size was observed both *in vitro* and *in vivo*. We found that small, newly budded cells tended to have larger capsules than their heavily pigmented mother cells and that cell size correlated positively with pigmentation in a melanizing culture. When a culture of cells with large capsules was provided with dopamine, a significant decrease in capsule thickness was observed within the first 6 h of melanization that was accompanied by a burst of polysaccharide release from cells. The growth and maintenance of large capsules was also impeded by exogenously provided melanin, but this effect could be reversed by addition of calcium to the culture media, which suggests calcium binding by melanin limits capsule size.

The cryptococcal capsule, a unique feature among fungal pathogens (31), is comprised predominantly of glucuronoxylomannon (GXM) that is synthesized intracellularly and exported in secretory vesicles for extracellular assembly (32). Cell wall melanin reduces the porosity of the cell wall (22) and slows vesicular transport (14), so we expected capsule growth for pigmented cells to be limited by decreased delivery of polysaccharide to the cell exterior. Remarkably, quantification of polysaccharide in culture supernatants by GXM ELISA revealed a higher concentration for melanized compared to non-melanized cells. These results argue that even if less polysaccharide is exported due to slowed vesicular traffic in melanized cells, so little was incorporated into the capsule that the net result is an increased abundance of shed polysaccharide for melanized compared to non-melanized cells.

It was possible to recapitulate inhibition of capsule growth and maintenance by providing melanin exogenously in a transwell system. This observation argues against a mechanism of melanin-induced modulation of the cell wall affecting polysaccharide attachment and instead suggests that melanin may be depleting a soluble factor required for capsule growth or assembly, such as an extracellular peptide that has been implicated in regulating virulence (33). Activated charcoal, which binds a wide range of organic and inorganic molecules, failed to reduce capsule growth. A notable difference between activated charcoal and melanin is that the former poorly adsorbs metals (34), while the latter has a high affinity for metal ions (25). The binding affinity of *Sepia* melanin for calcium measured by inductively coupled plasma mass spectrometry is 1.1 mmol/g (35), with an association constant determined by isothermal titration calorimetry of approximately 3.3 10^3^ M^-1^ (36) and we confirmed that cryptococcal melanin also binds calcium. Calcium binding by melanin is biologically relevant as melanin has been implicated in maintaining calcium homeostasis in human melanocytes (37). Since calcium is required by *C. neoformans* to form divalent crosslinks between polysaccharide chains during capsule assembly (26), we postulated that calcium binding to melanin may decrease the availability of calcium to negatively impact capsule growth. Our results support this hypothesis as calcium-supplemented culture media or pre-incubation of melanin with calcium diminished the inhibitory effect of melanin on capsule growth. Calcium binding to fungal melanin was recently shown to prevent phagocytosis of *Aspergillus* by disrupting calcium-calmodulin signaling (38) and our study has identified an additional mechanism by which calcium binding to melanin is able to modulate fungal virulence. We note that calcium supplementation did not fully restore the capsular size to that observed in the absence of melanin but divalent metal ion binding to the N-acetylglucosamine residues of capsular polysaccharide has an optimal stoichiometry of 1:2 such that capsule enlargement can be inhibited by too little or too much calcium (26). It is also possible that in addition to calcium, melanin is sequestering other components needed for capsule assembly. We also observed that melanization was associated with larger cells. Although we do not have an explanation for this observation, we note that the deposition of melanin in concentric layers (39) is dependent on cell wall flexibility (40), and this could require a larger volume to physically accommodate pigment granules circumferentially in the cell wall.

Cryptococcal virulence factors have been studied extensively for their singular contribution to pathogenic outcomes, but only recently have we recognized the potential for individual factors to interact in ways that may influence such outcomes (14). We have identified an inhibitory role for melanin in polysaccharide capsule growth and maintenance that has implications during infection as cryptococcal cells have been shown to become melanized in host tissue (41, 42). The abundance of melanin precursors in brain tissue is likely to favor melanization of disseminated *C. neoformans* cells and this may at least partially explain why the average cryptococcal capsule size in brain tissue is smaller than in lung tissue (43). Capsule enlargement has been observed to occur very soon after infection (44), but comparison of cells recovered from mouse lungs 24 h post-infection revealed significantly smaller capsules in tissue that had been infected with melanized compared to non-melanized *C. neoformans,* arguing that melanin is also able to inhibit capsule growth *in vivo*. A hallmark feature of cryptococcal infections is the accumulation of polysaccharide in host tissues and serum where it interferes with numerous immune processes, including inhibiting T-cell proliferation and interfering with secretion of proinflammatory cytokines (18). As we observed an initial burst of polysaccharide release during the first 6 h of melanization, deployment of this virulence factor during infection has the potential to increase the concentration of shed polysaccharide in tissues, thereby exacerbating its deleterious effects on host immunity.

Capsule assembly, although not fully understood, proceeds through direct anchoring of polysaccharides to *α*-1,3 glycan and chitin components of the cell wall (45, 46) followed by outward growth mediated by divalent metal ion crosslinking between polysaccharide chains (26). The resulting structure is non-uniform in its radial density, and consists of a denser inner capsule surrounded by a more sparse outer capsule (16). Comparison of capsule thicknesses before and after sonication showed a large decrease for non-melanized cells from 3 µm to 1 µm whereas melanized cells that already had a small average capsule thickness of 1.2 µm decreased only marginally to 1 µm. We quantified the volume and mass of capsules removed from non-melanized and melanized cells and found a 20-fold greater volume but only 2-fold greater mass for non-melanized cells compared to melanized cells. These results suggest that unlike non-melanized cells that have both inner and outer capsule layers, pigmented cells produce primarily a dense, inner capsule. We note that the diameter of polysaccharide molecules released from melanized cells were larger than those from non-melanized cells and attribute this to the fact that the former is capsular material whereas the latter is exopolysaccharide, which have different physical properties (47). The outer capsule can be removed by sonication, radiation, dimethyl sulfoxide or EDTA (23, 26, 48) but the inner capsule is resistant to these treatments and is tightly attached to the cell wall. Hence, the finding that melanized cells retain the inner capsule but lose their outer capsule provides a new tool for studying these structures. As calcium binding by melanin was observed to interfere with outer capsule growth, our results are consistent with a model wherein formation of the inner capsule layer is calcium-independent while assembly of the outer layer requires the formation of calcium bridges (Fig. 7).

**Fig. 7.**
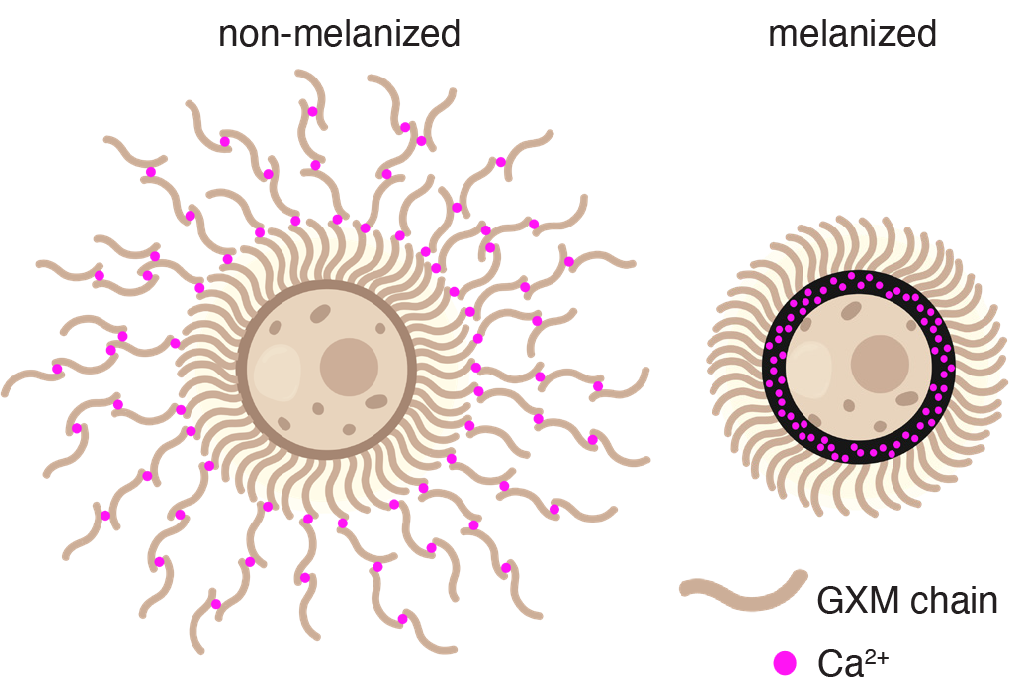
Model of calcium-dependent outer capsule assembly. Both non-melanized (left) and melanized (right) cells form dense inner capsules whereas growth of the calcium-dependent outer capsule that occurs readily in non-melanized cells is impeded by calcium binding to melanin. Illustration created with BioRender.com.

### Materials and Methods

***C. neoformans* cell culture.** Wild-type *Cryptococcus neoformans* serotype A strain KN99*α* (Fungal Genetics Stock Center, 2007 Lodge library) cells were recovered from frozen 50% glycerol stocks by growth at 30°C in yeast extract-peptone-dextrose (YPD) broth (BD Difco, 242820) before being sub-cultured into defined media for experiments. Capsule enlargement was induced by growth at 30°C in minimal media (MM) consisting of 29.4 mM K2HPO4, 10 mM MgSO4, 13 mM glycine, 15 mM D-glucose, and 3 µM thiamine at pH 5.5 in the absence or presence of 1 mM dopamine hydrochloride (Millipore Sigma, H8502).

### Capsule and pigment measurements

PBS-washed cells were mixed 1:1 with India ink (BD, 261194), wet mounted onto Fisherbrand Superfrost slides, and examined using an Olympus AX70 microscope under phase contrast with a 40X objective or under bright field illumination with a 100X oil-immersion objective. Images were captured using QCapture-Pro 6.0 software and a Retiga 1300 digital charge-coupled device camera. Cell body and total capsule plus cell body radius measurements in pixels were made from 100X images using the QCA Matlab script (49) or manually using Adobe Photoshop and measurements were converted to micrometers using the conversion factor of 0.0645 µm/pixel. Capsule thickness was calculated by subtracting the cell body radius from the total radius of capsule plus cell body. Pigment measurements were made by quantifying the mean gray value (MGV) of cell bodies in 100X images using ImageJ2 software. Mouse lung tissue analyzed in this study was recovered from frozen samples collected previously after infection of three animals with macrophages carrying either non-melanized or melanized *C. neoformans* (14). Pigmentation measurements plotted against capsule thickness or body radius were analyzed by simple linear regression and Pearson correlation using GraphPad Prism software.

### Melanization assay

Following growth in MM for 2 d at 30°C to induce capsule enlargement, cells were washed with PBS and then sonicated on ice at 18 W for 30 s using a Model 505 Sonic Dismembrator (Fisher Scientific). Samples taken pre- and post-sonication from three biological replicates were suspended in India ink and imaged for capsule measurements as described above. Sonicated and untreated cells were suspended at a density of 5 *×* 10^7^ cells/mL in PBS supplemented with 1 mM dopamine and incubated at 30°C. After 1, 2, 3, and 4 h, melanizing cells were transferred to wells of a 48-well plate and photographed using a 12-megapixel camera. Pigment intensities for each well were quantified using ImageJ2 software, plotted as MGV against time, and fit using simple linear regression. Pigmentation rates were adjusted for relative colony forming units (CFU) determined at each time point by plating diluted cultures on Sabouraud Dextrose (SAB) agar plates.

### Quantification of capsule size and polysaccharide released during melanization

Following capsule enlargement in MM, cells were washed with PBS, sub-cultured in triplicate at a cell density of 4 x 10^7^ cells/mL in dopamine-supplemented MM, and incubated at 30°C. Culture samples taken at 0, 6, 12, and 24 h were centrifuged at 2500 *g* for 5 min. Cell pellets were washed with PBS and suspended in India ink to obtain images used to derive capsule thickness measurements (as described above). Culture supernatants collected from 6, 12, and 24 h samples were passed through a 0.8 µm pore SFCA syringe filter (Corning, 431221) and quantified for polysaccharide content by GXM sandwich ELISA as described previously (50). Briefly, wells of high bind microplates (Corning, 9018) were coated with 1 µg/mL goat anti-mouse IgM in PBS, blocked using PBS with 1% BSA, and then incubated with 2 µg/mL anti-GXM antibody IgM-2D10. Diluted filtered supernatants and a 2.5µg/mL EPS standard were dispensed into the first row of the microplate and two-fold serially diluted into the remaining 7 rows. Captured GXM was detected by incubation with 5 µg/mL anti-GXM IgG- 18B7 followed by 1 µg/mL alkaline phosphate-labeled anti-IgG and finally 1 mg/mL phosphate substrate (Millipore Sigma, S0942) in a buffer comprised of 50 mM Na2CO3 and 1 mM MgCl2 at pH 9.8. Absorbance at 405 nm (A405) was measured using a Spectramax iD5 plate reader (Molecular Devices) and GXM content in supernatants was extrapolated from the linear equation derived from plotting A405 against concentration for the EPS standard. CFU quantified as described above were used to calculate the rate of GXM polysaccharide released per 10^8^ cells/h.

### Comparison of extracellular polysaccharide released from non-melanized and enriched melanized cultures

A heterogeneous culture of *C. neoformans* melanized for 2 d in 1 mM dopamine-supplemented MM was enriched for heavily melanized cells using density gradient centrifugation. Cells were layered onto 3 mL 70% stock isotonic Percoll prepared as described previously (15) and centrifuged at 40,000 rpm for 30 min in a TLA 100.3 fixed-angle rotor at 25°C. The layer of pale cells with large capsules at the interface of the dark band of melanized cells were discarded and the dark layer of heavily pigmented cells was transferred to a new tube. Three biological replicate cultures of enriched melanized cells and non-melanized control cells were washed with PBS and then sub-cultured at a density of 1 *×* 10^7^ cells/mL in MM with or without 1 mM dopamine, respectively. Following incubation for 4 h at 30°C, culture samples were diluted and plated on SAB agar plates to determine CFU. The remaining cells were pelleted by centrifugation at 3500 *g* for 5 min and supernatants were passed through a 0.8 µm syringe filter. Shed polysaccharide was quantified by GXM ELISA and release rates per 10^8^ cells/h were calculated.

### Quantification of volume and mass of polysaccharide released by sonication

Triplicate cultures of non-melanized and melanized cells (enriched for heavily pigmented cells as described above) were washed into PBS and sonicated at a cell density of 5 *×*10^7^ cells/mL for 30 s at 18W. Samples taken pre- and post-sonication were imaged in India ink and capsule and cell body radius measurements were made as described above and used to derive volumes using the equation V = 4/3*π*r^3^. Capsular volume measurements were calculated by subtracting the cell body volume from the total volume of capsule plus cell body. Sonicated cells were pelleted by centrifugation at 3500 *g* for 5 min, supernatants were passed through a 0.8 µm syringe filter, and the mass of polysaccharide was quantified by GXM ELISA and corrected for relative CFU quantified for each sonicated culture.

### Dynamic light scattering of polysaccharide released by sonication

Samples of polysaccharide collected by sonication of three replicate non-melanized and enriched melanized cultures, as described above, were dispensed into a 220-1600 nm UVette (Eppendorf, 952010051) and dynamic light scattering was performed using a Zeta Potential Analyzer (Brookhaven Instruments). ZetaPlus particle sizing software was used to determine multimodel size distributions and average particle sizes from 10 sequential 1 min measurements for each sample.

### Capsule regrowth following sonication

Following sonication (as described above), enriched melanized and non-melanized cells were sub-cultured in triplicate at a cell density of 1 *×* 10^7^ cells/mL in MM with and without 1 mM dopamine, respectively. Culture samples taken at 0, 4, and 17 h were washed with PBS, suspended in India ink, and imaged for capsule thickness measurements as described above. GXM ELISA was used to quantify the amount of polysaccharide in 0.8µm-filtered supernatants collected at 4 and 17 h time intervals and CFU counts were used to normalize measurements for 10^8^ cells.

### Transwell capsule growth and maintenance assays

For capsule growth analysis, cells from a YPD culture were washed with PBS and then sub-cultured in triplicate in polyethylene terephthalate 0.4 µm permeable cell culture inserts (Celltreat) at a density of 2 *×*10^6^ cells/mL. Outer wells contained either MM or MM supplemented with 3 mg/mL synthetic melanin (Millipore Sigma, M0418), 5 mg/mL cryptococcal melanin ghosts prepared from a dopamine-supplemented *C. neoformans* culture as described previously (24), or 5 mg/mL activated charcoal (Millipore Sigma, 31616). Following incubation at 30°C with shaking at 200 rpm for 48 h, PBS-washed cells were suspended in India ink and imaged for capsule measurements (as described above). A similar procedure was used to examine the effect of melanin on capsule maintenance except cells were cultured in MM at 30°C for 2 d to induce capsule enlargement before being sub-cultured into the transwell system with MM or MM supplemented with 0.5, 1, or 2 mg/mL synthetic melanin. Synthetic melanin was suspended in water or a 1M solution of CaCl2 for 3 d before being washed three times with water and lyophilized. Capsule growth inhibition was compared in triplicate transwells containing MM supplemented with 2 mg/mL the calcium-treated and control. Capsule growth was also assayed in the absence or presence of 1-1.5 mg/mL synthetic melanin in MM or magnesium-free MM with or without the addition of 0.1 mM calcium chloride as indicated in figure legends.

### Calcium binding to *C. neoformans* melanin ghosts and synthetic melanin

Suspensions of melanin ghosts in duplicate or synthetic melanin in triplicate were prepared at a concentration of 1 mg/mL in calcium chloride solutions ranging from 5 mM to 0.5 M. Following equilibration for 1 h at room temperature, each suspension was centrifuged at 5,000 *g* for 5 min and the supernatant was passed through a 0.22 µm syringe filter. The concentration of calcium in the unbound fraction was quantified using a LAQUAtwin Ca-11 calcium meter (Horiba) and these values were subtracted from the original concentration to determine the amount of bound calcium. A saturation binding curve was plotted and fit by the equation Y = Bmax *×* X/(Kd + X) using GraphPad Prism software.

## Acknowledgments

We are grateful for support from the Bloomberg Distinguished Professor Summer Program (A.Z.L.) and the National Institutes of Health, grant number R01-HL059842 (A.C.).

